# Noradrenergic and Dopaminergic Neural Correlates of Trait Anxiety: Unveiling the Impact of Maladaptive Emotion Regulation

**DOI:** 10.1101/2024.07.23.604801

**Authors:** Michal Rafal Zareba, Paula Ariño-Braña, Maria Picó-Pérez, Maya Visser

## Abstract

**Background:** Maladaptive emotion regulation plays a crucial role in the development and maintenance of elevated anxiety levels, both in patients and in individuals with subclinical symptomatology. While pharmacological treatments for anxiety target the emotion dysregulation through dopaminergic, noradrenergic and serotonergic systems, little is known about the underlying mechanisms. Therefore, the current study depicts the association of these neuromodulatory systems’ resting-state functioning with trait-anxiety, investigating the role of maladaptive emotion regulation.

**Methods:** Amplitude of low-frequency fluctuations (ALFF), fractional amplitude of low-frequency fluctuations (fALFF), and whole-brain resting-state functional connectivity (rs-FC) were obtained from the ventral tegmental area (VTA), locus coeruleus (LC) and dorsal raphe, and correlated with trait-anxiety and self-reported maladaptive emotion regulation (N = 60).

**Results:** Trait-anxiety was positively associated with LC’s fALFF and negatively with VTA’s whole-brain rs-FC with the left inferior parietal lobule (L-IPL) and the left superior frontal gyrus (L-SFG). Maladaptive emotion regulation was negatively associated with VTA’s rs-FC with these regions, with trait-anxiety fully mediating this association. VTA connectivity with the frontal region, but not parietal, positively predicted its amplitude of neural oscillations, an effect that was paralleled by stronger frontal dopaminergic innervation.

**Limitations:** Use of external molecular datasets and need for replication in patients.

**Conclusions:** Noradrenergic and dopaminergic systems appear to contribute differently to subclinical anxiety. While noradrenaline likely acts through a more general mechanism, the dopaminergic dysconnectivity with the frontoparietal control network may act as one of the mechanisms of maladaptive emotion regulation, informing the models on the disorder development.

**Highlights:**

- Trait-anxiety shows a positive association with the activity of locus coeruleus
- Trait-anxiety and emotional regulation are linked to VTA-frontoparietal connectivity
- Trait-anxiety fully mediates emotion regulation’s effect on VTA’s connectivity
- Strength of connectivity with VTA is positively linked to the frontal ALFF and fALFF

## 1. Introduction

Anxiety disorders constitute a prevalent category of mental health problems that affect individuals’ overall well-being, accompanied by physical symptoms such as fatigue, tiredness, and irritability (Wu et al., 2024). Despite the variability across anxiety disorders (Purves et al., 2020), all patients encounter similar challenges in regulating their emotions, often resorting to maladaptive strategies such as avoidance, rumination, catastrophizing, and self-blame, instead of adaptive ones like cognitive reappraisal (Garnefski et al., 2002; Rodríguez-Rey et al., 2024). Importantly, this characteristic is believed to both give rise to and reinforce the anxiety symptoms (Rodriguez-Rey et al., 2024), and extends beyond clinical diagnosis. Subclinical symptomatology or high trait-anxiety have similarly been associated with impaired emotion regulation processes (Cécillon et al., 2024, Brehl et al., 2021). Owing to the meta-analytic efforts, the brain basis of these maladaptive processes and negative emotional biases in anxiety have been identified. Both diagnosed individuals and those with high levels of trait-anxiety display hyper-reactivity in affective brain regions such as the amygdala and insula, which may contribute to heightened fear experiences (Etkin & Wager, 2007). Furthermore, brain areas involved in cognitive reappraisal, such as the cingulate and prefrontal cortices, as well as the angular gyri, exhibit hypoactivation, indicating a failure in cognitive control of negative emotions (Picó-Pérez et al., 2017).

As these brain areas are densely innervated by the brainstem monoamine neuromodulators — dopamine, norepinephrine and serotonin — drugs targeting these systems are commonly prescribed as pharmaceutical therapies for anxiety disorders. Dysregulation of the dopaminergic activity in the ventral tegmental area (VTA) has been linked to anxiety symptoms across species (Peng et al., 2021; Zweifel et al., 2011; Cha et al., 2014: Zalachoras et al., 2022). Similar results have been found for the impaired noradrenergic functions of locus coeruleus (LC; Bouras et al., 2023, Grueschow et al., 2020) and serotonergic neurotransmission in dorsal raphe (DR; Hale et al., 2012; Chen et al., 2020). However, despite the apparent involvement of these midbrain areas in maladaptive emotion regulation and anxiety, the specific mechanisms by which their projections affect this symptomatology remain unclear (Abdul et al., 2022; Payet et al., 2023), which might hinder our understanding of the present therapies’ processes and efficacy (Li et al., 2020; Cao et al., 2021).

One particularly useful tool for characterising the functioning of the brain comes in the form of resting-state functional magnetic resonance imaging (rs-fMRI). While most studies on particular diagnoses rely on task-based fMRI with paradigms involving emotion processing or regulation, task and resting-state scans may not be entirely separate measures for psychopathology. Emerging data indicates that the patterns represented by both types of measures overlap significantly, accounting for more than 80% of the variance (Elliott et al., 2019). One benefit of using rs-fMRI for studying mental health is that it allows non-invasive and effortless data acquisition from both healthy and clinical populations, who might have difficulties performing cognitive tasks inside the scanner (Lv et al., 2019). Its clinical applicability is further encouraged by straightforwardness of data analysis (Ge & Dou, 2022), and the recent advances in the identification of potential biomarkers using such data (Zugman et al., 2023; Yang et al., 2023).

One way to better understand the brain regions involved in the described phenomena is to investigate their functioning not only in clinical patients, but also in highly anxious people who do not meet the criteria for a clinical diagnosis. The current understanding of psychopathology on a dimensional scale is limited as research is still frequently conducted comparing healthy individuals with clinically diagnosed subjects (Besteher et al., 2017). To challenge this, initiatives such as the Research Domain Criteria (Kozak and Cuthbert, 2016) proposed that narrower psychological constructs such as cognition or emotion might be more susceptible to biological analysis than relying on traditional diagnostic categories such as anxiety or depression. Following this idea, we understand psychopathology as a continuum, which requires studying the underlying neural substrates within this spectrum. Thus, investigating subclinical populations with high trait-anxiety could help us get a more complete comprehension of the link between the brain bases of emotion regulation and anxiety (Laeger et al., 2012), as well as provide us with important implications for treatment. Furthermore, since subclinical populations are not typically under any psychopharmacological treatment, targeting these populations allows us to explore the neural correlates of the psychological process of interest without the modulatory effect of medication being at play.

In this regard, the existing neuroimaging literature provides insightful perspectives on the neural correlates of specific anxiety disorders, particularly within fronto-limbic and fronto-parietal connectivity (Kim et al., 2011; Hein & Ruiz, 2022). However, there is a lack of direct investigation in humans into how trait-anxiety is related to the neural activity of the midbrain regions, as well as their connectivity with the aforementioned cortical circuitry. Furthermore, despite the role of maladaptive emotion regulation in the genesis of anxiety disorders (Garnefski et al., 2002; Rodriguez-Rey et al., 2024), currently it is unknown whether this phenomenon is in fact fully responsible for the neural patterns associated with increased anxiety levels. A more complete understanding of these processes is crucial given that currently prescribed pharmacological treatments primarily exert their therapeutic effects through these neuromodulatory systems.

Therefore, we set out to investigate the functioning of VTA, LC and DR in relation to individual variation in trait-anxiety. We measured the spontaneous resting-state neural activity occurring in these brain regions using the amplitude of low-frequency fluctuations (ALFF), as well as a variation of this technique called the fractional amplitude of low-frequency fluctuations (fALFF). Considering the evidence from animal research and single positron emission tomography (PET) studies in humans, we expected to observe positive associations of trait-anxiety with the ALFF and fALFF of LC and DR (Bouras et al., 2023; Hale et al., 2012; Chen et al., 2020). In the case of the dopaminergic system, both increased and decreased VTA activity has been reported in high trait-anxiety (Zalachoras et al., 2022, Berry et al., 2019), and as such our investigation was meant to weigh additional arguments into the ongoing debate.

We additionally hypothesised that trait-anxiety would be associated with resting-state functional connectivity between our regions of interests (ROIs) and frontoparietal regions relevant for emotional processing and regulation. We expected to observe both negative and positive correlations, taking into account the recent evidence from rodents showing that manipulating distinct connections of the same neuromodulatory region can exert opposite effects on behaviour (Ren et al., 2018). Furthermore, given the well-established relationship that the inefficient emotion regulation leads to higher anxiety (Rodriguez-Rey et al., 2024), we hypothesised that the use of maladaptive emotion regulation strategies would be associated with overlapping neural correlates with trait-anxiety, and that trait-anxiety would act as a mediator of this relationship. Providing such novel evidence for the neuromodulatory systems’ contributions would greatly benefit the theories on the emergence of anxiety symptomatology in humans, as well as potential treatment mechanisms of pharmacological and psychotherapeutic first-line treatments.

Following this principle, our study also aimed to assess the neurogenetic underpinnings of the neuromodulatory systems in relation to their impact on neural connectivity as influenced by trait-anxiety levels. Firstly, we wanted to investigate whether the ALFF and fALFF of the regions identified as significant in the connectivity analyses would be associated with the strength of their connectivity with the respective brainstem seeds. Such an association is indeed to be expected if the brainstem neurotransmission exerts a modulatory influence on the downstream brain areas. Secondly, as a post-hoc analysis, we aimed to examine the expression patterns of neurotransmitter receptors and transporters in these brain regions to uncover the molecular bases of these observations. By integrating neuroimaging findings with neurogenetic data, we hope to provide a more comprehensive understanding of the neural mechanisms involved in trait-anxiety, thereby contributing to a multi-level analysis of brain function and its molecular determinants.

## 2. Methods

### 2.1. Participants

The sample for this study was selected from the “Max Planck Institute Mind-Brain-Body Dataset” (Babayan et al., 2019), an openly accessible dataset that provides anonymised MRI and behavioural data in compliance with the Declaration of Helsinki’s ethical standards. Participants were required to meet the specific inclusion criteria: 20-35 years old, being right-handed, possessing at least a secondary-level education, and reporting neither current medication use nor a history of neurological, psychiatric, or substance use disorders. These preliminary requirements were met by 94 individuals. Given the lack of complete behavioural data for 5 individuals and excessive head motion in 29 subjects (i.e. maximal volume displacement larger than the voxel size), the final sample consisted of 60 participants (20 females). The age of participants was available only in the form of 5 year brackets (e.g. 20-25) as the authors of the dataset wanted to ensure the complete anonymity of their participants, nevertheless, this does not interfere with the goals of the current study. The demographic summary of the final sample is provided in Table 1.

**Table 1.** Demographic and behavioural data.

|  |  |  |
| --- | --- | --- |
| **Sex/gender (N, %)** | **Males** | 40 (66.67%) |
|  | **Females** | 20 (33.33%) |
| **Age range (N, %)** | **20-25** | 30 (50%) |
|  | **25-30** | 25 (41.67%) |
|  | **30-35** | 5 (8.33%) |
| **Trait-anxiety - STAI (Mean ± SD)** | 36.95 ± 8.04 | |
| **Maladaptive emotion regulation - CERQ (Mean ± SD)** | 14.58 ± 5.68 | |
| **Self-blame - CERQ subscale (Mean ± SD)** | 4.27 ± 2.02 | |
| **Rumination - CERQ subscale (Mean ± SD)** | 5.83 ± 2.74 | |
| **Catastrophizing - CERQ subscale (Mean ± SD)** | 2.12 ± 1.84 | |
| **Blaming others - CERQ subscale (Mean ± SD)** | 2.53 ± 1.90 | |

Abbreviations: STAI, State-Trait-Anxiety Inventory (Spielberger et al., 1970); CERQ, Cognitive Emotion Regulation Questionnaire (Garnefski et al., 2001); SD, standard deviation.

### 2.2. Behavioural data

Trait-anxiety was assessed with the short version of the State-Trait-Anxiety Inventory (STAI; Spielberger et al., 1970; Laux et al., 1981). The scale assesses trait-anxiety levels through 20 items rated on a Likert scale from 1 (almost never) to 4 (nearly always). The participants additionally completed the Cognitive Emotion Regulation Questionnaire (CERQ; Garnefski et al., 2001; Loch et al., 2011), which probed the use of various adaptive and maladaptive cognitive approaches to emotion regulation through 27 Likert-scale items marked from 0 (almost never) to 4 (almost always). The maladaptive emotion regulation strategies score was obtained by summing up the subscores from the self-blame, other-blame, rumination and catastrophizing categories.

### 2.3. MRI data acquisition

Acquisitions were performed on a 3 Tesla scanner (MAGNETOM Verio, Siemens Healthcare GmbH, Erlangen, Germany) equipped with a 32-channel head coil. Structural images were obtained with an MPRAGE sequence (176 sagittal slices; 1 mm isotropic voxel size; TR = 5000 ms; TE = 2.92 ms; flip angle 1/flip angle 2 = 4°/5°; GRAPPA acceleration factor 3). For the T2*-weighted resting-state functional sequence, participants were asked to remain awake with their eyes concentrated on a low-contrast fixation cross. The data was acquired using a gradient echo planar multiband imaging sequence (64 axial slices acquired in interleaved order; 2.3 mm isotropic voxel size; TR = 1400 ms; TE = 30 ms; flip angle = 69°; multiband acceleration factor = 4). 657 volumes were obtained, giving the total scan duration of 15 mins 30 s.

### 2.4. MRI data preprocessing and (f)ALFF computation

The first 5 volumes for the fMRI data were discarded to ensure signal stabilisation. Such functional images and original anatomical data were preprocessed using the fMRIPrep 23.0.0. (Esteban et al., 2018), which is based on Nipype 1.8.5. (Gorgolewski et al., 2011). A comprehensive overview of the preprocessing steps is available in the Supplementary Material. Shortly, the anatomical images were skull-stripped, segmented and normalised to the MNI space. The functional data underwent motion, susceptibility distortion and slice-timing correction, coregistration to the anatomical image and normalisation. The remaining steps of the functional preprocessing were performed in AFNI (Cox, 1996). The fMRI volumes were smoothed with a 4 mm Gaussian filter, and were subsequently detrended, denoised and band-pass filtered in the 0.01-0.1 frequency range in one step. The denoising regressors were taken from the output of fMRIprep, and they included 6 motion parameters and their derivatives, and component-based physiological regressors (aCompCor; Behzadi et al., 2007) for cerebrospinal fluid (CSF) explaining half of the CSF signal variance. For fALFF, polynomial detrending was not applied, following recent recommendations (Woletz et al., 2019).

Using AFNI software (Cox, 1996), ALFF was calculated as the total signal power within the 0.01-0.1 Hz frequency range, whereas fALFF was measured as a ratio of the signal power within the aforementioned range divided by the signal power of the entire frequency spectrum. Both metrics were mean-normalised within each individual prior to the group analyses. While ALFF shows higher test-retest reliability than fALFF in grey matter regions (Zuo et al., 2010), fALFF is known to be less sensitive to the physiological noise (Zou et al., 2008). As such, we decided to use both metrics in our investigation, which would allow us to compare the results between them.

### 2.5. Regions of interest delineation and signal extraction

The ROIs for DR, bilateral LC and VTA were delineated in the MNI space using publicly available atlases (Murty et al., 2014; Beliveau et al., 2015; Betts et al., 2017). Compared to demarking these areas in each brain, the current approach is much more feasible for the potential clinical use due to the lack of need for radiological expertise. After resampling the ROI templates to the resolution of the functional images, DR consisted of 9 voxels, while bilateral LC had 15 voxels in total. As the original VTA ROI was much larger than DR and LC (54 voxels), we aimed to reduce its size in order to exclude the potential differences in ALFF, fALFF and connectivity as a consequence of spatial extent discrepancies. The probabilistic VTA ROI was thus first thresholded at 0.999 probability, and subsequently a 1.5 voxel size (3.45 mm) sphere was drawn at the mass centre of the thresholded image (x = 0, y = -18, z = -14) to be used as an inclusive mask. The resulting VTA ROI consisted of 14 voxels. The location of ROIs in the MNI space are depicted in the Supplementary Figure 1.

For each ROI in each participant, we extracted the mean ALFF and fALFF values. Additionally, we created the seed time courses of the blood-oxygen-level-dependent (BOLD) signal by averaging the time-series from all voxels in each region. Subsequently, we computed the Pearson correlations between these mean time courses and all the other voxels in the brain, and normalised the distribution of the correlation coefficients using Fisher’s z transform.

### 2.6. Statistical Analysis

#### 2.6.1. Group-level analyses for ALFF and fALLF data

Apart from the whole-brain imaging analyses, all the statistical procedures were performed in R (version 2023.06.0; R Core Team, 2023). The associations of the amplitudes of resting-state neural activity with trait-anxiety were tested using linear regression models. ALFF or fALFF of each ROI was treated as the dependent variable, while trait-anxiety, sex/gender and age were used as regressors. The correction for multiple comparisons was achieved using the false-discovery rate (FDR < 0.05). As the aim of our study was to see whether maladaptive emotion regulation would be responsible for the emergence of neural patterns associated with trait-anxiety, the linear models using this characteristic as the regressor were run only for the neural data highlighted in the anxiety-related analyses.

#### 2.6.2. Group-level analyses for functional connectivity data

Firstly, one sample t-tests of the whole-brain functional connectivity patterns of each ROI were performed using AFNI’s 3dttest++ program to check that the group-averaged connectivity maps were consistent with previous literature and ensure proper seed delineation. In turn, the effects of trait-anxiety on the whole-brain functional connectivity patterns of the seeds were examined with the use of AFNI’s 3dMVM program (Chen et al., 2014). Similarly to the ALFF and fALFF analyses, sex/gender and age were included as covariates. Correction for multiple comparisons was achieved with a cluster-level family-wise error rate correction (FWE < 0.05) following initial voxel-level thresholding at the level of p < 0.005.

Subsequently, we extracted the mean functional connectivity values of the significant clusters from the previous analyses, and tested whether the use of maladaptive emotion regulation strategies was also their significant predictor using linear regression. The effects of age and sex/gender were already included in the whole-brain models and thus controlled. The results were FDR-corrected for multiple comparisons.

#### 2.6.3. Mediation analysis

To examine whether trait-anxiety mediates the relationship between the use of maladaptive emotion regulation strategies and the associated neural markers, we used R’s mediation package (Tingley et al., 2014). The mediation model included maladaptive emotion regulation strategies as the independent variable, trait-anxiety as the mediator, and the neural markers as the dependent variable. This analysis was performed only for those neural markers that were found to be significantly associated with both trait-anxiety and maladaptive emotion regulation.

#### 2.6.4. Associations between the connectivity strength and neural signal amplitude

Lastly, we ran linear regression models to test whether the strength of the connectivity between the brainstem ROIs and the significant clusters from the group-level trait-anxiety analyses could predict the latter’s ALFF and fALFF.

## 3. Results

### 3.1. Trait-anxiety is positively related to fALFF in locus coeruleus

In line with our hypothesis, we observed a positive association between the levels of trait-anxiety and fALFF in the LC (t = 2.93, p_FDR_ = 0.015), indicating that the higher the trait-anxiety levels, the higher the spontaneous activity in this area (Figure 1). We did not find, however, any other significant results for the amplitude measures. Similarly, the use of maladaptive emotion regulation strategies was not related to LC fALLF (t = 0.496, p = 0.622).

**Figure 1.**
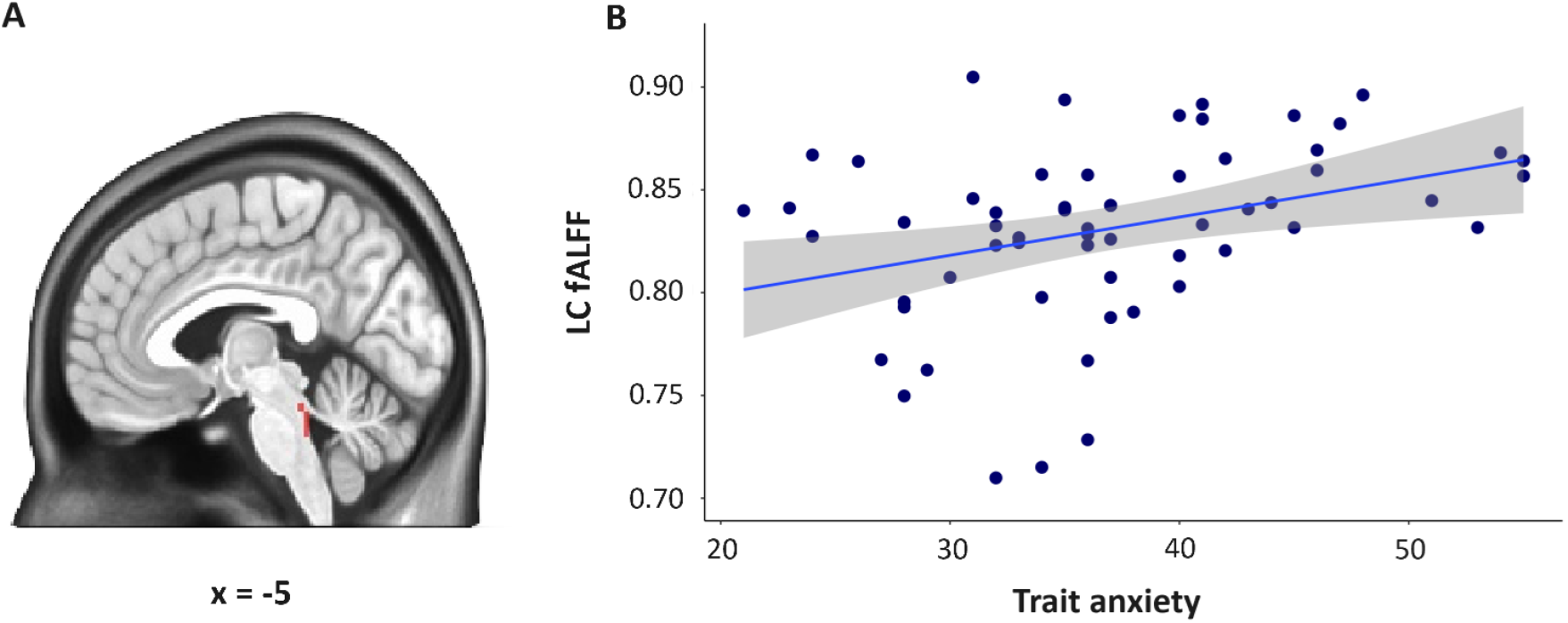
Association between trait-anxiety and fALFF in the locus coeruleus (LC). A. LC ROI, highlighted in red. B. Plot illustrating the positive association between trait-anxiety levels and LC fALFF. The higher the trait-anxiety levels, the higher the spontaneous low frequency activity in this area.

### 3.2. Trait-anxiety fully mediates the association between maladaptive emotion regulation and ventral tegmental area connectivity

One sample t-tests revealed that the functional connectivity patterns of LC, VTA and DR were in line with those reported in the literature, thus confirming their correct delineation (Beliveau et al., 2015; Murty et al., 2014; Zareba et al., 2022). The results are presented in Supplementary Figures 2-4.

As for the main analysis, a significant negative effect of trait-anxiety emerged on the functional connectivity of VTA with two cortical regions (Figure 2 and Table 2). Higher trait-anxiety levels were associated with lower connectivity between the VTA and the left inferior parietal lobule (L-IPL; r = -0.56) and the left superior frontal gyrus (L-SFG; r = -0.57). The functional connectivity of VTA with L-IPL (t = -2.11, p_FDR_ = 0.044) and L-SFG (t = -2.06, p_FDR_ = 0.044) was also related to maladaptive emotion regulation. As for LC and DR, there were no statistically significant effects of trait-anxiety on the whole-brain functional connectivity. The unthresholded statistical maps for all seeds are published alongside the manuscript.

**Table 2.** Significant effects of trait-anxiety on the whole-brain resting-state functional connectivity of the ventral tegmental area.

| MNI coordinates (x, y, z) | Area | Voxels | F-stat | Pearson correlation |
| --- | --- | --- | --- | --- |
| -51, -71, 43 | L inferior parietal lobule | 135 | 20.35 | -0.56 |
| -2, 24, 46 | L superior frontal gyrus | 73 | 21.78 | -0.57 |

**Figure 2.**
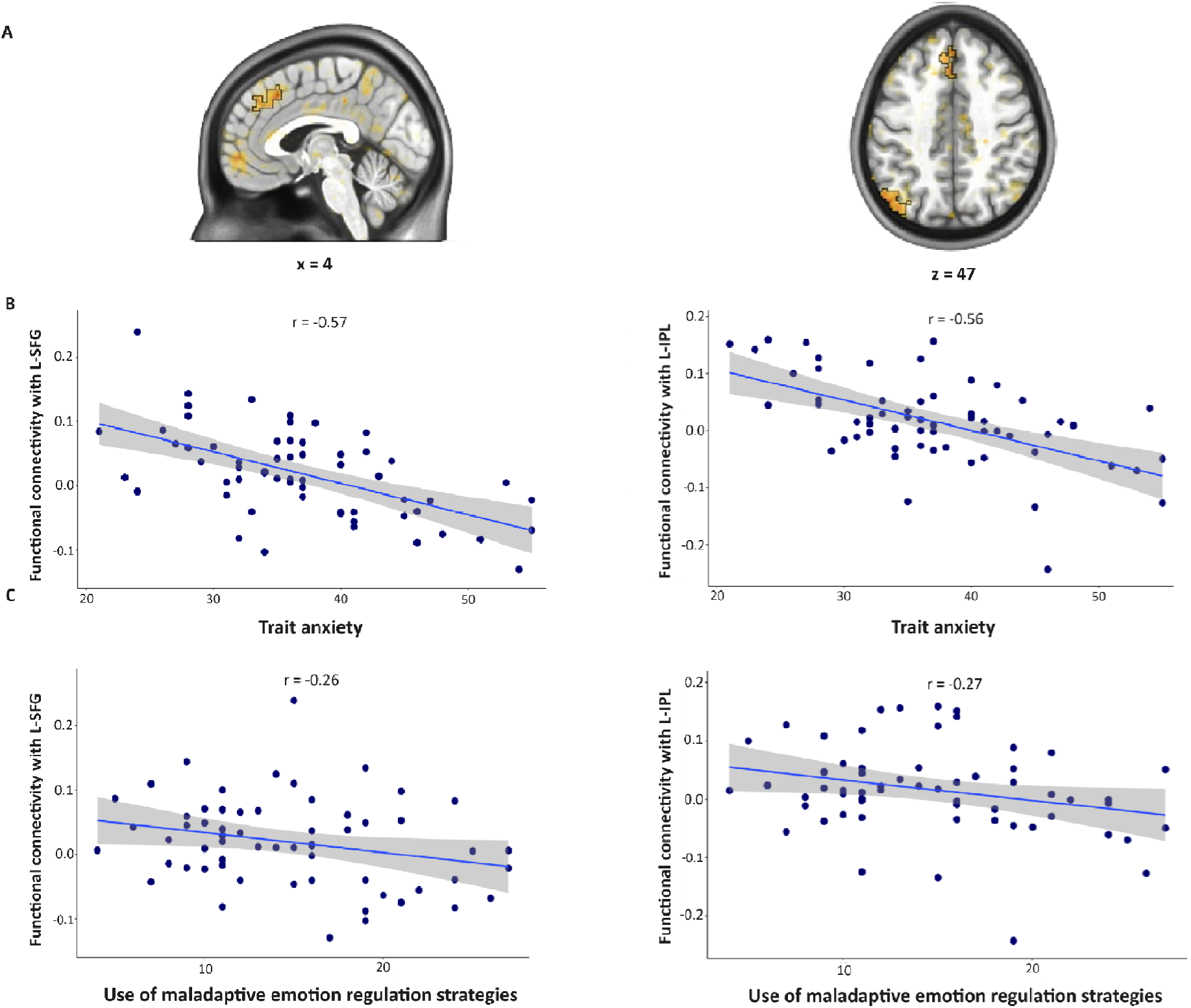
Associations of trait-anxiety and maladaptive emotion regulation with ventral tegmental area’s (VTA) resting-state functional connectivity (rs-FC). A. Regions showing reduced rs-FC with VTA in association with trait-anxiety levels and maladaptive emotion regulation strategies (outlined in black). B. Plots illustrating the significant negative association of trait-anxiety with VTA’s rs-FC with these regions. C. Plots illustrating the associations of maladaptive emotion regulation strategies with VTA’s rs-FC with these regions. In both cases, higher trait-anxiety levels and the use of maladaptive regulation strategies were associated with lower rs-FC between the VTA and these two structures.

Considering maladaptive emotion regulation was significantly associated with trait-anxiety (t = 4.39, p < 0.001), and both measures were linked with VTA’s functional connectivity, we tested a mediation model in which the relationship between maladaptive emotion regulation and the connectivity patterns would be mediated by trait-anxiety. Indeed, trait-anxiety was found to be fully mediating the associations for both brain regions, thus confirming the study’s hypotheses (Figure 3).

**Figure 3.**
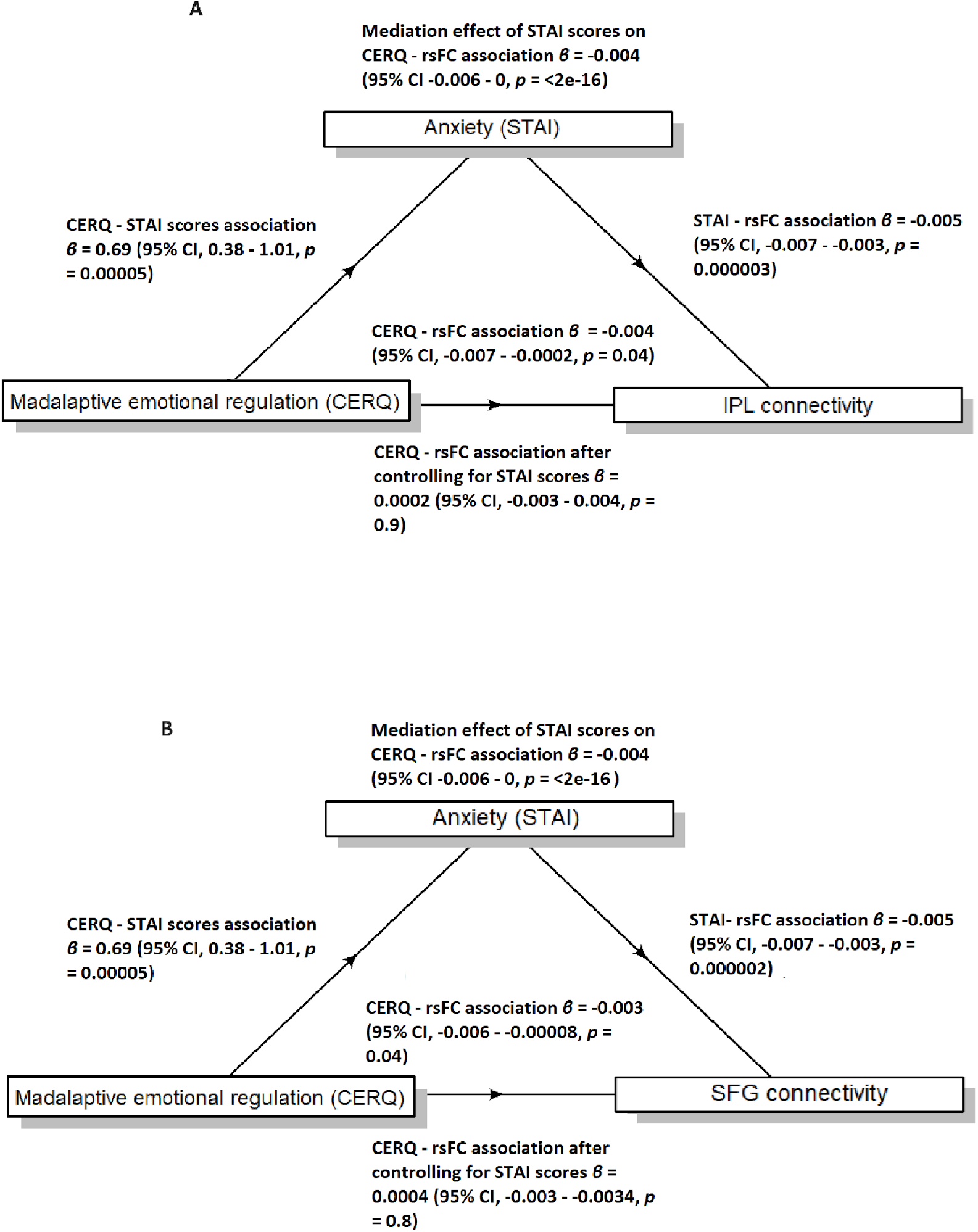
Trait-anxiety acts as a full mediator of the effect of the use of maladaptive emotion regulation strategies on ventral tegmental area (VTA) - frontoparietal resting-state functional connectivity (rs-FC). A. The mediation plot for the left inferior parietal lobule (L-IPL). B. The mediation plot for the left superior frontal gyrus (L-SFG).

### 3.3. Associations between the connectivity strength and neural signal amplitude, and their regional molecular characteristics

The functional connectivity between VTA and L-SFG significantly predicted ALFF (t = 3.50; p < 0.001) and fALFF (t = 4.71; p < 0.001) of the frontal region. However, such associations were not observed neither for ALFF (t = -1.16; p = 0.252), nor fALFF (t = -1.17; p = 0.246) of the L-IPL, which probed us to investigate whether the two regions differed in their molecular characteristics related to the dopaminergic system.

As such, we used the external dopamine D1 and D2 receptor gene expression maps (Gryglewski et al., 2018) and PET-based dopamine transporter (DAT) maps (Dukart et al., 2018; Hansen et al., 2022). The maps were mean-normalised and resampled to the fMRI data resolution. Following the recent findings that D1/D2 receptor ratio was linked to the effects of methylphenidate administration on fALFF across the brain, we decided to use this measure instead of D1 and D2 expression patterns themselves (Manza et al., 2022). The values for D1/D2 ratio and DAT were extracted for each voxel from the significant clusters from the connectivity analyses and compared between regions using t-tests. These analyses revealed that L-SFG had profoundly higher DAT density compared to L-IPL (t = 15.16; p < 0.001; Cohen’s d = 4.25). In turn, L-IPL was found to have significantly higher D1/D2 receptor ratio (t = 3.52; p < 0.001; Cohen’s d = 0.60).

## 4. Discussion

Our study was aimed at increasing the understanding of the mechanisms behind the contributions of noradrenergic, dopaminergic and serotonergic systems to the individual differences in trait-anxiety. In line with our hypotheses, the first major finding was the significant positive effect of trait-anxiety on LC’s fALFF. LC, the major brain source of noradrenaline, is known to play a crucial role in attention, vigilance, and arousal, particularly under stress conditions (Daviu et al., 2019; Regelink, 2020).

A body of works points towards LC’s crucial role in the development of anxiety disorders, as well as its hyperresponsiveness in already highly anxious individuals (Inoue & Koyama, 2012; Etkin & Wager, 2007; Qin et al., 2021; Ji et al., 2023; Garnefski et al., 2018). Animal studies suggest that stress, which is an important factor in the genesis of anxiety, primarily biases LC towards high tonic activity and diminishes its differential responses to discrete stimuli (McCall et al., 2015). This pattern in turn contributes to basolateral amygdala’s hypersensitivity to emotional stimuli (McCall et al., 2017; Brehl et al., 2020), and disrupts the functioning of the cognitive control circuitry within the prefrontal cortex (Lapiz & Morilak, 2006). Furthermore, a recent study in humans has linked trait-anxiety to the LC’s structural connectivity with dorsal attention, salience, and frontoparietal attention networks, underlining the important role of the noradrenergic innervation in both internally- and externally-guided cognition, as well as their integration (Liebe et al., 2022).

Given how broad the spectrum of processes affected by noradrenergic neurotransmission is, as well as the absence of specific functional connectivity patterns related to trait-anxiety in the current and previous results (Liebe et al., 2022), we believe that increased noradrenergic tone may have a generic mechanism of action, affecting the excitability of multiple brain areas to a similar extent. Despite the evidence for negative associations of adaptive emotional regulation and increased LC-controlled basal sympathetic nervous system activity (Golkar et al., 2014), we do not report a parallel association with our data. Similar bidirectional links have also been reported for the more transient noradrenergic activity (Raio et al., 2013; Gross, 1998), indicating the possible interplay between the two modes of neurotransmission in regards to emotion regulation, which may be difficult to grasp with non-invasive imaging techniques. Nevertheless, the positive association between trait-anxiety and LC’s fALFF reported here indicates that rs-fMRI could still be a promising tool for the assessment of noradrenergic activity. If replicated in a clinical cohort, this neural signature could potentially aid in the search for neuroimaging biomarkers of anxiety disorders.

Nevertheless, contrary to our hypotheses, we did not find any significant results regarding the association of trait-anxiety with DR’s and VTA’s signal amplitude. One potential explanation for it could be the functional heterogeneity of the serotonergic and dopaminergic neurons. It has been demonstrated that activation of serotonergic neurons can have both anxiogenic and anxiolytic effects depending on their target structure (Ren et al., 2018), whereas distinct populations of dopaminergic neurons encode motivational value, being excited by reward stimuli and inhibited by aversive events, and motivational salience, being excited by both rewards and aversive stimuli (Bromberg-Martin et al., 2010). As such, it is possible that the link between trait-anxiety and the activity of serotonergic and dopaminergic neurons may depend on their functional profile, meaning that within one brain region we may encounter neurons with both upregulated and downregulated activity. As in the case of ALFF and fALLF we observe the net effect within a particular area, this heterogeneity may limit our ability to detect such findings.

The second main finding of our study was that trait-anxiety and the use of maladaptive emotion regulation strategies were both associated with lower functional connectivity of VTA with L-IPL and L-SFG, and that the relationship between emotion regulation and these connectivity patterns was fully mediated by trait-anxiety. These results are in line with previous evidence supporting the role of maladaptive emotional regulation in the development and maintenance of elevated levels of anxiety (Rodriguez-Rey et al., 2024), however, they highlight for the first time the direct contributions of the dopaminergic system. Furthermore, our work provides an important insight that this process likely occurs through cortical connection-specific mechanisms, as compared to the generic noradrenergic neuromodulation.

L-IPL and L-SFG are both the key components of the lateral frontoparietal network responsible for executive functions, and as such they naturally play a pivotal role in cognitive emotion regulation (Menon, 2011; Martín-Signes et al., 2021). L-SFG, which maps onto the presupplementary motor area (preSMA), is associated with the use of language and memory processing needed to reformulate mental representations (Wang et al., 2018; Jackson et al., 2021), which could be viewed as an instance of action selection (Wager et al., 2008). In turn, the parietal regions are involved in maintaining attention in the reappraised thoughts and emotions (Wang et al., 2018; Jackson et al., 2021). Recent meta-analyses have found that in anxiety patients both regions were hypoactive during cognitive reappraisal (Picó-Pérez et al., 2017), and that their increased baseline activation positively predicted the psychotherapy outcome (Picó-Pérez et al., 2023). While our study is cross-sectional in its nature, it is a well established fact that dopaminergic neurotransmission affects the excitability of cortical neurons on a time scale longer than that of individual action potentials, thus altering the dynamics of interregional cortical communication (Lavin & Grace, 2001). Therefore, with the prominent role of dopamine in action selection and attentional processes (Rowe et al., 2010; Bensmann et al., 2020; Elton et al., 2021), the presented results suggest the decreased connectivity with the dopaminergic centres as one of the mechanisms for maladaptive emotion regulation. Although the current research uses the resting-state paradigm, there is growing evidence that the resting-state neural patterns closely align with task-based neural dynamics, and might even have a role in shaping task-based activity (Feng et al., 2022; Elliott et al., 2019). Thus, our results indeed might be associated with the performance of individuals with high trait-anxiety in emotionally demanding situations.

While more studies are required in this line of research to fully confirm this interpretation, especially task-based, the present work and other reports provide preliminary support for these claims. In studies deploying acute pharmacological challenges to decrease or increase the brain dopamine levels, the cortical functional connectivity of VTA usually follows the neuromodulator’s trajectory (Kline et al., 2016; Grimm et al., 2020; Elton et al., 2021). In one case, lowering dopaminergic availability simultaneously reduced attentional biases towards rewarding stimuli (Elton et al., 2021). In turn, increasing dopamine levels was also reported to increase fALFF in the associative cortices, including the regions reported in our study, and decrease the signal amplitude in the sensorimotor areas, a differential characteristic that was linked to higher D1/D2 ratio in the former regions (Manza et al., 2022). In the current work, the strength of VTA connectivity with L-SFG was positively related to the latter’s ALFF and fALFF, indicating that resting-state neuroimaging can potentially be a viable tool for studying the neuromodulatory effects of dopaminergic neurotransmission even in the absence of pharmacological challenges.

Nevertheless, as mentioned in the results, the lack of such findings for L-IPL probed us towards investigating the differences in molecular characteristics between the two brain regions by using freely available maps of DAT availability (Dukart et al., 2018; Hansen et al., 2022), and dopamine D1 and D2 receptor gene expression (Gryglewski et al., 2018). While L-IPL was found to have a higher D1/D2 receptor ratio than L-SFG (Cohen’s d = 0.60), the latter region had a far more abundant strength of the dopaminergic innervation, as evident with the DAT levels (Cohen’s d = 4.25). As such, these results suggest that the neuromodulatory effects of dopamine may be detectable in the resting-state paradigm only for areas that receive relatively strong dopaminergic input, as compared to acute pharmacological challenges (Manza et al., 2022). Nonetheless, given the fact that research on the dopamine system is focused mainly on regions with dense dopaminergic innervation, our work hints at a new and easily accessible avenue for the use of resting-state in this line of studies.

The use of external molecular data poses, however, one of the limitations of the study. Nevertheless, this preliminary step opens up potential new research directions and warrants the works exploring the usability of such an approach with the use of subject-specific PET datasets. Secondly, given the subclinical characteristics of our sample, caution needs to be taken generalising these findings until they are replicated in clinical groups. As such, it would be of interest to see whether the markers of subclinical anxiety identified here could potentially predict the progression of anxiety symptomatology to a clinical severity. Last but not least, while our findings are consistent with task-based studies, the inherent complexities of interpreting resting-state data should also be noted out.

In conclusion, the current work provides novel insights into the relationship of trait-anxiety with the activity and functional connectivity of human midbrain regions. In more detail, it suggests differential modes of contribution of the noradrenergic and dopaminergic systems, which appear to act, respectively, through generic and connection-specific mechanisms. The presented results furthermore underline the role of the maladaptive emotion regulation in the genesis of anxiety, for the first time linking it to the dysconnectivity of the dopaminergic centres with the lateral frontoparietal control network. Last but not least, the positive association between the VTA connectivity and the amplitude of neural oscillations in the frontal cortex indicates that dopamine might shape the functional characteristics of the downstream regions even during rest, suggesting a possible mechanism for the sustenance of maladaptive emotion regulation and subclinical anxiety.

## Data availability

The unthresholded statistical maps generated for this study will be made openly available on the NeuroVault platform upon manuscript acceptance.

## CRediT author statement

MRZ: Conceptualization, Formal analysis, Visualization, Writing - Original Draft. PAB: Conceptualization, Formal analysis, Visualization, Writing - Original Draft. MPP: Conceptualization, Writing - Review & Editing. MV: Conceptualization, Supervision, Writing - Review & Editing.

## Ethical statement

This work was performed using an anonymised openly available dataset (Babayan et al., 2019) and thus was not subjected to ethical committee approval.

## References

Abdul, M., Yan, H. Q., Zhao, W. N., Lyu, X. B., Xu, Z., Yu, X. L., … & Cao, J. L. (2022). VTA-NAc glutaminergic projection involves in the regulation of pain and pain-related anxiety. Frontiers in Molecular Neuroscience, 15, 1083671.

Babayan, A., Erbey, M., Kumral, D., Reinelt, J. D., Reiter, A. M., Röbbig, J., … & Villringer, A. (2019). A mind-brain-body dataset of MRI, EEG, cognition, emotion, and peripheral physiology in young and old adults. Scientific data, 6(1), 1–21. Behzadi, Yashar, Khaled Restom, Joy Liau, and Thomas T. Liu. 2007. “A Component Based Noise Correction Method (CompCor) for BOLD and Perfusion Based fMRI.” NeuroImage 37 (1): 90–101. 10.1016/j.neuroimage.2007.04.042.

Beliveau, V., Svarer, C., Frokjaer, V. G., Knudsen, G. M., Greve, D. N., & Fisher, P. M. (2015). Functional connectivity of the dorsal and median raphe nuclei at rest. NeuroImage, 116, 187–195. 10.1016/j.neuroimage.2015.04.065

Bensmann, W., Zink, N., Arning, L., Beste, C., & Stock, A. K. (2020). Dopamine D1, but not D2, signaling protects mental representations from distracting bottom-up influences. Neuroimage, 204, 116243.

Berry, A. S., White, R. L., Furman, D. J., Naskolnakorn, J. R., Shah, V. D., D’Esposito, M., & Jagust, W. J. (2019). Dopaminergic mechanisms underlying normal variation in trait-anxiety. Journal of Neuroscience, 39(14), 2735–2744.

Besteher, B., Gaser, C., Langbein, K., Dietzek, M., Sauer, H., & Nenadić, I. (2017). Effects of subclinical depression, anxiety and somatization on brain structure in healthy subjects. Journal of affective disorders, 215, 111–117.

Betts, M. J., Cardenas-Blanco, A., Kanowski, M., Jessen, F., & Düzel, E. (2017). In vivo MRI assessment of the human locus coeruleus along its rostrocaudal extent in young and older adults. NeuroImage, 163, 150–159. 10.1016/j.neuroimage.2017.09.042

Bouras, N. N., Mack, N. R., & Gao, W. J. (2023). Prefrontal modulation of anxiety through a lens of noradrenergic signaling. Frontiers in Systems Neuroscience, 17, 1173326.

Brehl, A. K., Schene, A., Kohn, N., & Fernández, G. (2021). Maladaptive emotion regulation strategies in a vulnerable population predict increased anxiety during the Covid-19 pandemic: A pseudo-prospective study. Journal of Affective Disorders Reports, 4, 100113.

Brehl, A. K., Kohn, N., Schene, A. H., & Fernández, G. (2020). A mechanistic model for individualised treatment of anxiety disorders based on predictive neural biomarkers. Psychological Medicine, 50(5), 727–736.

Bromberg-Martin, E. S., Matsumoto, M., & Hikosaka, O. (2010). Dopamine in motivational control: rewarding, aversive, and alerting. Neuron, 68(5), 815–834. 10.1016/j.neuron.2010.11.022

Cao, J., Huang, Y., Hodges, S. A., Meshberg, N., & Kong, J. (2021). Identify potential neuroimaging-based scalp acupuncture and neuromodulation targets for anxiety. Brain Science Advances, 7(2), 97–111.

Cécillon, F. X., Mermillod, M., Leys, C., Lachaux, J. P., Le Vigouroux, S., & Shankland, R. (2024). trait-anxiety, Emotion Regulation, and Metacognitive Beliefs: An Observational Study Incorporating Separate Network and Correlation Analyses to Examine Associations with Executive Functions and Academic Achievement. Children, 11(1), 123.

Cha, J., Carlson, J. M., DeDora, D. J., Greenberg, T., Proudfit, G. H., & Mujica-Parodi, L. R. (2014). Hyper-reactive human ventral tegmental area and aberrant mesocorticolimbic connectivity in overgeneralization of fear in generalized anxiety disorder. Journal of Neuroscience, 34(17), 5855–5860.

Chen, G., Adleman, N. E., Saad, Z. S., Leibenluft, E., & Cox, R. W. (2014). Applications of multivariate modeling to neuroimaging group analysis: a comprehensive alternative to univariate general linear model. Neuroimage, 99, 571–588. Chen, Chenyi et al. “An integrative analysis of 5HTT-mediated mechanism of hyperactivity to non-threatening voices.” Communications biology vol. 3, 1 113. 10 Mar. 2020, doi:10.1038/s42003-020-0850-3

Cox, R. W. (1996). AFNI: software for analysis and visualization of functional magnetic resonance neuroimages. Computers and Biomedical research, 29(3), 162–173. Daviu, N., Bruchas, M. R., Moghaddam, B., Sandi, C., & Beyeler, A. (2019). Neurobiological links between stress and anxiety. Neurobiology of stress, 11, 100191.

Dukart, J., Holiga, Š., Chatham, C., Hawkins, P., Forsyth, A., McMillan, R., Myers, J., Lingford-Hughes, A. R., Nutt, D. J., Merlo-Pich, E., Risterucci, C., Boak, L., Umbricht, D., Schobel, S., Liu, T., Mehta, M. A., Zelaya, F. O., Williams, S. C., Brown, G., Paulus, M., … Sambataro, F. (2018). Cerebral blood flow predicts differential neurotransmitter activity. Scientific reports, 8(1), 4074. 10.1038/s41598-018-22444-0

Elliott, M. L., Knodt, A. R., Cooke, M., Kim, M. J., Melzer, T. R., Keenan, R., … & Hariri, A. R. (2019). General functional connectivity: Shared features of resting-state and task fMRI drive reliable and heritable individual differences in functional brain networks. Neuroimage, 189, 516–532.

Elton, A., Faulkner, M. L., Robinson, D. L., & Boettiger, C. A. (2021). Acute depletion of dopamine precursors in the human brain: effects on functional connectivity and alcohol attentional bias. Neuropsychopharmacology : oficial

Esteban, O., Markiewicz, C. J., Blair, R. W., Moodie, C. A., Isik, A. I., Erramuzpe, A., … & Gorgolewski, K. J. (2019). fMRIPrep: a robust preprocessing pipeline for functional MRI. Nature methods, 16(1), 111–116.

Etkin, A., & Wager, T. D. (2007). Functional neuroimaging of anxiety: a meta-analysis of emotional processing in PTSD, social anxiety disorder, and specific phobia. American journal of Psychiatry, 164(10), 1476–1488. publication of the American College of Neuropsychopharmacology, 46(8), 1421–1431. 10.1038/s41386-021-00993-9

Feng, Z. J., Deng, X. P., Zhao, N., Jin, J., Yue, J., Hu, Y. S., … & Wang, J. (2022). Resting-state fMRI functional connectivity strength predicts local activity change in the dorsal cingulate cortex: A multi-target focused rTMS study. Cerebral Cortex, 32(13), 2773–2784.

Garnefski, N., Kraaij, V., & Spinhoven, P. (2001). Negative life events, cognitive emotion regulation and emotional problems. Personality and Individual differences, 30(8), 1311–1327.

Garnefski, N., Legerstee, J., Kraaij, V., van Den Kommer, T., & Teerds, J. A. N. (2002). Cognitive coping strategies and symptoms of depression and anxiety: A comparison between adolescents and adults. Journal of adolescence, 25(6), 603–611.

Garnefski, N., & Kraaij, V. (2018). Specificity of relations between adolescents’ cognitive emotion regulation strategies and symptoms of depression and anxiety. Cognition and emotion, 32(7), 1401–1408.

Golkar, A., Johansson, E., Kasahara, M., Osika, W., Perski, A., & Savic, I. (2014). The influence of work-related chronic stress on the regulation of emotion and on functional connectivity in the brain. PloS one, 9(9), e104550.

Gorgolewski, K., Burns, C. D., Madison, C., Clark, D., Halchenko, Y. O., Waskom, M. L., & Ghosh, S. S. (2011). Nipype: a flexible, lightweight and extensible neuroimaging data processing framework in python. Frontiers in neuroinformatics, 5, 13.

Grimm, O., Kopfer, V., Küpper-Tetzel, L., Deppert, V., Kuhn, M., de Greck, M., & Reif, A. (2020). Amisulpride and l-DOPA modulate subcortical brain nuclei connectivity in resting-state pharmacologic magnetic resonance imaging. Human brain mapping, 41(7), 1806–1818. 10.1002/hbm.24913

Grueschow, M., Kleim, B., & Ruff, C. C. (2020). Role of the locus coeruleus arousal system in cognitive control. Journal of Neuroendocrinology, 32(12), e12890.

Gryglewski, G., Seiger, R., James, G. M., Godbersen, G. M., Komorowski, A., Unterholzner, J., Michenthaler, P., Hahn, A., Wadsak, W., Mitterhauser, M., Kasper, S., & Lanzenberger, R. (2018). Spatial analysis and high resolution mapping of the human whole-brain transcriptome for integrative analysis in neuroimaging. NeuroImage, 176, 259–267. 10.1016/j.neuroimage.2018.04.068

Hansen, J. Y., Shafiei, G., Markello, R. D., Smart, K., Cox, S. M. L., Nørgaard, M., Beliveau, V., Wu, Y., Gallezot, J. D., Aumont, É., Servaes, S., Scala, S. G., DuBois, J. M., Wainstein, G., Bezgin, G., Funck, T., Schmitz, T. W., Spreng, R. N., Galovic, M., Koepp, M. J., … Misic, B. (2022). Mapping neurotransmitter systems to the structural and functional organization of the human neocortex. Nature neuroscience, 25(11), 1569–1581. 10.1038/s41593-022-01186-3

Hale, M. W., Shekhar, A., & Lowry, C. A. (2012). Stress-related serotonergic systems: implications for symptomatology of anxiety and affective disorders. Cellular and molecular neurobiology, 32, 695–708.

Hein, T. P., & Ruiz, M. H. (2022). State anxiety alters the neural oscillatory correlates of predictions and prediction errors during reward-based learning. Neuroimage, 249, 118895.

Inoue T, Koyama T. [Anxiety disorder]. Brain Nerve. 2012 Feb;64(2):131–8. Japanese. PMID: 22308258.

Jackson, R. L. (2021). The neural correlates of semantic control revisited. NeuroImage, 224, 117444.

Ji, Q., Li, S. J., Zhao, J. B., Xiong, Y., Du, X. H., Wang, C. X., … & Zhu, Z. R. (2023). Genetic and neural mechanisms of sleep disorders in children with autism spectrum disorder: a review. Frontiers in Psychiatry, 14, 1079683.

Kline, R. L., Zhang, S., Farr, O. M., Hu, S., Zaborszky, L., Samanez-Larkin, G. R., & Li, C. S. (2016). The Effects of Methylphenidate on Resting-State Functional Connectivity of the Basal Nucleus of Meynert, Locus Coeruleus, and Ventral Tegmental Area in Healthy Adults. Frontiers in human neuroscience, 10, 149. 10.3389/fnhum.2016.00149

Kozak MJ, Cuthbert BN. The NIMH Research Domain Criteria Initiative: Background, Issues, and Pragmatics. Psychophysiology. 2016 Mar;53(3):286–97. doi: 10.1111/psyp.12518. PMID: 26877115.

Laeger, I., Dobel, C., Dannlowski, U., Kugel, H., Grotegerd, D., Kissler, J., … & Zwanzger, P. (2012). Amygdala responsiveness to emotional words is modulated by subclinical anxiety and depression. Behavioural Brain Research, 233(2), 508–516.

Lapiz, M. D. S., & Morilak, D. A. (2006). Noradrenergic modulation of cognitive function in rat medial prefrontal cortex as measured by attentional set shifting capability. Neuroscience, 137 (3), 1039–1049.

Laux, L., Glanzmann, P., Schaffner, P., & Spielberger, C. (1981). Das State-Trait-Angstinventar. Theoretische Grundlagen und Handanweisung. Weinheim: Beltz Test GmbH.

Lavin, A., & Grace, A. A. (2001). Stimulation of D1-type dopamine receptors enhances excitability in prefrontal cortical pyramidal neurons in a state-dependent manner. Neuroscience, 104 (2), 335–346. 10.1016/s0306-4522(01)00096-3

Li, J., Zhong, Y., Ma, Z., Wu, Y., Pang, M., Wang, C., et al. (2020). Emotion reactivity-related brain network analysis in generalized anxiety disorder: A task fMRI study. BMC Psychiatry, 20, 429. 10.1186/s12888-020-02831-6

Li, W., Yang, P., Ngetich, R., Zhang, J., Jin, Z., & Li, L. (2021). Differential involvement of frontoparietal network and insula cortex in emotion regulation. Neuropsychologia, 161, 107991. 10.1016/j.neuropsychologia.2021.107991

Liebe, T., Kaufmann, J., Hämmerer, D., Betts, M., & Walter, M. (2022). In vivo tractography of human locus coeruleus—relation to 7T resting state fMRI, psychological measures and single subject validity. Molecular Psychiatry, 27 (12), 4984–4993.

Loch, N., Hiller, W., & Witthöft, M. (2011). Der Cognitive Emotion Regulation Questionnaire (CERQ). Zeitschrift für Klinische Psychologie und Psychotherapie, 40 (2), 94–106.

Lv, H., Wang, Z., Tong, E., Williams, L. M., & Wintermark, M. (2018). Resting-State Functional MRI: Everything That Nonexperts Have Always Wanted to Know. American Journal of Neuroradiology. 10.3174/ajnr.A5527

Manza, P., Shokri-Kojori, E., Demiral, Ş.B., Wiers, C. E., Zhang, R., Giddens, N., … & Volkow, N. D. (2022). Cortical D1 and D2 dopamine receptor availability modulate methylphenidate-induced changes in brain activity and functional connectivity. Communications Biology, 5 (1), 514.

Martín-Signes, M., Cano-Melle, C., & Chica, A. B. (2021). Fronto-parietal networks underlie the interaction between executive control and conscious perception: evidence from TMS and DWI. Cortex, 134, 1–15. 10.1016/j.cortex.2020.09.027

McCall, J. G., Al-Hasani, R., Siuda, E. R., Hong, D. Y., Norris, A. J., Ford, C. P., & Bruchas, M. R. (2015). CRH Engagement of the Locus Coeruleus Noradrenergic System Mediates Stress-Induced Anxiety. Neuron, 87 (3), 605–620. doi: 10.1016/j.neuron.2015.07.002

McCall, J. G., Siuda, E. R., Bhatti, D. L., Lawson, L. A., McElligott, Z. A., Stuber, G. D., & Bruchas, M. R. (2017). Locus coeruleus to basolateral amygdala noradrenergic projections promote anxiety-like behavior. eLife, 6, e18247.

Menon, V. (2011). Large-scale brain networks and psychopathology: a unifying triple network model. Trends in Cognitive Sciences, 15 (10), 483–506.

Murty, V. P., Shermohammed, M., Smith, D. V., Carter, R. M., Huettel, S. A., & Adcock, R. A. (2014). Resting state networks distinguish human ventral tegmental area from substantia nigra. NeuroImage, 100, 580–589. 10.1016/j.neuroimage.2014.06.047

Payet, J. M., Stevens, L., Russo, A. M., Jaehne, E. J., van den Buuse, M., Kent, S., … & Hale, M. W. (2023). The Role of Dorsal Raphe Nucleus Serotonergic Systems in Emotional Learning and Memory in Male BALB/c Mice. Neuroscience, 534, 1–15.

Peng, B., Xu, Q., Liu, J., Guo, S., Borgland, S. L., & Liu, S. (2021). Corticosterone attenuates reward-seeking behavior and increases anxiety via D2 receptor signaling in ventral tegmental area dopamine neurons. Journal of Neuroscience, 41(7), 1566–1581.

Picó-Pérez, M., Fullana, M. A., Albajes-Eizagirre, A., Vega, D., Marco-Pallarés, J., Vilar, A., … & Soriano-Mas, C. (2023). Neural predictors of cognitive-behavior therapy outcome in anxiety-related disorders: a meta-analysis of task-based fMRI studies. Psychological Medicine, 53(8), 3387–3395.

Picó-Pérez, M., Radua, J., Steward, T., Menchón, J. M., & Soriano-Mas, C. (2017). Emotion regulation in mood and anxiety disorders: a meta-analysis of fMRI cognitive reappraisal studies. Progress in Neuro-Psychopharmacology and Biological Psychiatry, 79, 96–104. Wu, J., Li, X., Zhang, Q., Li, J., Cui, R., & Li, X. (2024). Differential effects of intra-RMTg infusions of pilocarpine or 4-DAMP on regulating depression-and anxiety-like behaviors. Behavioural Brain Research, 462, 114833.

Purves, K. L., Coleman, J. R., Meier, S. M., Rayner, C., Davis, K. A., Cheesman, R., … & Eley, T. C. (2020). A major role for common genetic variation in anxiety disorders. Molecular psychiatry, 25(12), 3292–3303.

Qin, X., Liu, X. X., Wang, Y., Wang, D., Song, Y., Zou, J. X., … & Zhang, W. H. (2021). Early life stress induces anxiety-like behavior during adulthood through dysregulation of neuronal plasticity in the basolateral amygdala. Life Sciences, 285, 119959.

R Core Team. (2023). R: A language and environment for statistical computing. R Foundation for Statistical Computing, Vienna, Austria. URL https://www.R-project.org/.

Regelink, M. (2020). Locus Coeruleus Drive in Relation to Trait-Anxiety and Behavioral Performance during Stress.

Ren, J., Friedmann, D., Xiong, J., Liu, C. D., Ferguson, B. R., Weerakkody, T., DeLoach, K. E., Ran, C., Pun, A., Sun, Y., Weissbourd, B., Neve, R. L., Huguenard, J. R., Horowitz, L. F., & Luo, L. (2018). Anatomically defined and functionally distinct dorsal raphe serotonin sub-systems. Cell, 175 (2), 472-487.e20. doi:10.1016/j.cell.2018.07.043

Rodríguez-Rey, R., Guerra Corral, M., Collazo-Castiñeira, P., Collado, S., Caro-Carretero, R., Cantizano, A., & Garrido-Hernansaiz, H. (2024). Predictors of mental health in healthcare workers during the COVID-19 pandemic: The role of experiential avoidance, emotion regulation and resilience. Journal of Advanced Nursing.

Rowe, J. B., Hughes, L. E., Barker, R. A., & Owen, A. M. (2010). Dynamic causal modelling of effective connectivity from fMRI: Are results reproducible and sensitive to Parkinson’s disease and its treatment? NeuroImage, 52 (3), 1015–1026. 10.1016/j.neuroimage.2009.12.080

Spielberger, C. D. (1970). Manual for the State-Trait Anxiety Inventory (self-evaluation questionnaire).

Tingley, D., Yamamoto, T., Hirose, K., Keele, L., & Imai, K. (2014). mediation: R Package for Causal Mediation Analysis. In Journal of Statistical Software (Vol. 59, Issue 5, pp. 1–38). http://www.jstatsoft.org/v59/i05/

Wager, T. D., Davidson, M. L., Hughes, B. L., Lindquist, M. A., & Ochsner, K. N. (2008). Neural mechanisms of emotion regulation: evidence for two independent prefrontal-subcortical pathways. Neuron, 59(6), 1037.

Wang, H. Y., Zhang, X. X., Si, C. P., Xu, Y., Liu, Q., Bian, H. T., … & Yan, Z. R. (2018). Prefrontoparietal dysfunction during emotion regulation in anxiety disorder: a meta-analysis of functional magnetic resonance imaging studies. Neuropsychiatric disease and treatment, 1183–1198.

Woletz, M., Hoffmann, A., Tik, M., Sladky, R., Lanzenberger, R., Robinson, S., & Windischberger, C. (2019). Beware detrending: Optimal preprocessing pipeline for low-frequency fluctuation analysis. Human brain mapping, 40(5), 1571–1582. 10.1002/hbm.24468

Yang, B., Wang, X., Mo, J., Li, Z., Hu, W., Zhang, C., Zhao, B., Gao, D., Zhang, X., Zou, L., Zhao, X., Guo, Z., Zhang, J., & Zhang, K. (2023). The altered spontaneous neural activity in patients with Parkinson’s disease and its predictive value for the motor improvement of deep brain stimulation. NeuroImage. Clinical, 38, 103430. 10.1016/j.nicl.2023.103430

Zalachoras, I., Astori, S., Meijer, M., Grosse, J., Zanoletti, O., de Suduiraut, I. G., … & Sandi, C. (2022). Opposite effects of stress on effortful motivation in high and low anxiety are mediated by CRHR1 in the VTA. Science advances, 8(12), eabj9019.

Zareba, M. R., Furman, W., & Binder, M. (2022). Influence of age and cognitive performance on resting-state functional connectivity of dopaminergic and noradrenergic centers. Brain Research, 1796, 148082.

Zou, Q. H., Zhu, C. Z., Yang, Y., Zuo, X. N., Long, X. Y., Cao, Q. J., … & Zang, Y. F. (2008). An improved approach to detection of amplitude of low-frequency fluctuation (ALFF) for resting-state fMRI: fractional ALFF. Journal of neuroscience methods, 172(1), 137–141. Zugman, A., Jett, L., Antonacci, C., Winkler, A. M., & Pine, D. S. (2023). A Systematic Review and Meta-Analysis of Resting-state fMRI in Anxiety Disorders: Need for Data Sharing to Move the Field Forward. Journal of anxiety disorders, 102773.

Zuo, X. N., Di Martino, A., Kelly, C., Shehzad, Z. E., Gee, D. G., Klein, D. F., … & Milham, M. P. (2010). The oscillating brain: complex and reliable. Neuroimage, 49(2), 1432–1445. Zweifel, L. S., Fadok, J. P., Argilli, E., Garelick, M. G., Jones, G. L., Dickerson, T. M., … & Palmiter, R. D. (2011). Activation of dopamine neurons is critical for aversive conditioning and prevention of generalized anxiety. Nature neuroscience, 14(5), 620–626.

