## Supplementary Materials for "Noradrenergic and Dopaminergic Neural Correlates of Trait Anxiety: Unveiling the Impact of Maladaptive Emotion Regulation"

### 1. Details of the fMRIprep (version 23.0.0.) pipeline

Results included in this manuscript come from preprocessing performed using fMRIPrep 23.0.0 (Esteban, Markiewicz, et al. (2018); Esteban, Blair, et al. (2018); RRID:SCR_016216), which is based on Nipype 1.8.5 (K. Gorgolewski et al. (2011); K. J. Gorgolewski et al. (2018); RRID:SCR_002502).

#### 1.1. Preprocessing of B0 inhomogeneity mappings

A total of 1 fieldmaps were found available within the input BIDS structure for each subject. A B0-nonuniformity map (or fieldmap) was estimated based on two (or more) echo-planar imaging (EPI) references with topup (Andersson, Skare, and Ashburner (2003); FSL 6.0.5.1:57b01774).

#### 1.2. Anatomical data preprocessing

A total of 1 T1-weighted (T1w) images were found within the input BIDS dataset.The T1-weighted (T1w) image was corrected for intensity non-uniformity (INU) with N4BiasFieldCorrection (Tustison et al. 2010), distributed with ANTs 2.3.3 (Avants et al. 2008, RRID:SCR_004757), and used as T1w-reference throughout the workflow. The T1w-reference was then skull-stripped with a Nipype implementation of the antsBrainExtraction.sh workflow (from ANTs), using OASIS30ANTs as target template. Brain tissue segmentation of cerebrospinal fluid (CSF), white-matter (WM) and gray-matter (GM) was performed on the brain-extracted T1w using fast (FSL 6.0.5.1:57b01774, RRID:SCR_002823, Zhang, Brady, and Smith 2001). Volume-based spatial normalization to one standard space (MNI152NLin2009cAsym) was performed through nonlinear registration with antsRegistration (ANTs 2.3.3), using brain-extracted versions of both T1w reference and the T1w template. The following template was selected for spatial normalization: ICBM 152 Nonlinear Asymmetrical template version 2009c [Fonov et al. (2009), RRID:SCR_008796; TemplateFlow ID: MNI152NLin2009cAsym].

#### 1.3. Functional data preprocessing

For the 1 BOLD run found per subject, the following preprocessing was performed. First, a reference volume and its skull-stripped version were generated using a custom methodology of fMRIPrep. Head-motion parameters with respect to the BOLD reference (transformation matrices, and six corresponding rotation and translation parameters) are estimated before any spatiotemporal filtering using mcflirt (FSL 6.0.5.1:57b01774, Jenkinson et al. 2002). The estimated fieldmap was then aligned with rigid-registration to the target EPI (echo-planar imaging) reference run. The field coefficients were mapped on to the reference EPI using the transform. BOLD runs were slice-time corrected to 0.659s (0.5 of slice acquisition range 0s-1.32s) using 3dTshift from AFNI (Cox and Hyde 1997, RRID:SCR_005927). The BOLD reference was then co-registered to the T1w reference using mri_coreg (FreeSurfer) followed by flirt (FSL 6.0.5.1:57b01774, Jenkinson and Smith 2001) with the boundary-based registration (Greve and Fischl 2009) cost-function. Co-registration was configured with six degrees of freedom. Several confounding time-series were calculated based on the preprocessed BOLD: framewise displacement (FD), DVARS and three region-wise global signals. FD was computed using two formulations following Power (absolute sum of relative motions, Power et al. (2014)) and Jenkinson (relative root mean square displacement between affines, Jenkinson et al. (2002)). FD and DVARS are calculated for each functional run, both using their implementations in Nipype (following the definitions by Power et al. 2014). The three global signals are extracted within the CSF, the WM, and the whole-brain masks. Additionally, a set of physiological regressors were extracted to allow for component-based noise correction (CompCor, Behzadi et al. 2007). Principal components are estimated after high-pass filtering the preprocessed BOLD time-series (using a discrete cosine filter with 128s cut-off) for the two CompCor variants: temporal (tCompCor) and anatomical (aCompCor). tCompCor components are then calculated from the top 2% variable voxels within the brain mask. For aCompCor, three probabilistic masks (CSF, WM and combined CSF+WM) are generated in anatomical space. The implementation differs from that of Behzadi et al. in that instead of eroding the masks by 2 pixels on BOLD space, a mask of pixels that likely contain a volume fraction of GM is subtracted from the aCompCor masks. This mask is obtained by thresholding the corresponding partial volume map at 0.05, and it ensures components are not extracted from voxels containing a minimal fraction of GM. Finally, these masks are resampled into BOLD space and binarized by thresholding at 0.99 (as in the original implementation). Components are also calculated separately within the WM and CSF masks. For each CompCor decomposition, the k components with the largest singular values are retained, such that the retained components' time series are sufficient to explain 50 percent of variance across the nuisance mask (CSF, WM, combined, or temporal). The remaining components are dropped from consideration. The head-motion estimates calculated in the correction step were also placed within the corresponding confounds file. The confound time series derived from head motion estimates and global signals were expanded with the inclusion of temporal derivatives and quadratic terms for each (Satterthwaite et al. 2013). Frames that exceeded a threshold of 0.25 mm FD or 1.5 standardized DVARS were annotated as motion outliers. Additional nuisance timeseries are calculated by means of principal components analysis of the signal found within a thin band (crown) of voxels around the edge of the brain, as proposed by (Patriat, Reynolds, and Birn 2017). The BOLD time-series were resampled into standard space, generating a preprocessed BOLD run in MNI152NLin2009cAsym space. First, a reference volume and its skull-stripped version were generated using a custom methodology of fMRIPrep. All resamplings can be performed with a single interpolation step by composing all the pertinent transformations (i.e. head-motion transform matrices, susceptibility distortion correction when available, and co-registrations to anatomical and output spaces). Gridded (volumetric) resamplings were performed using antsApplyTransforms (ANTs), configured with Lanczos interpolation to minimize the smoothing effects of other kernels (Lanczos 1964). Non-gridded (surface) resamplings were performed using mri_vol2surf (FreeSurfer).

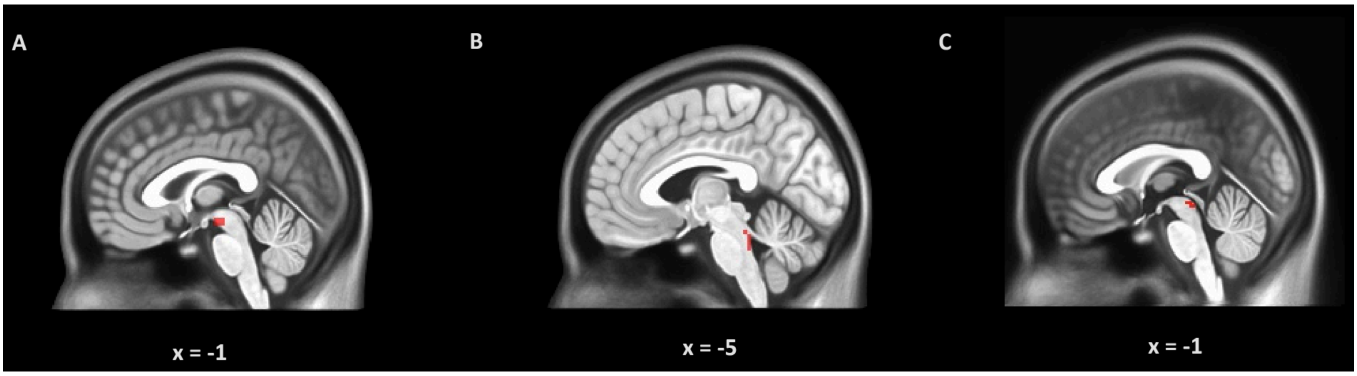


**Supplementary Figure 1.** The location of regions of interests (ROIs) in the MNI space. A. Ventral tegmental area (VTA). B. Locus coeruleus (LC). C. Dorsal raphe (DR).

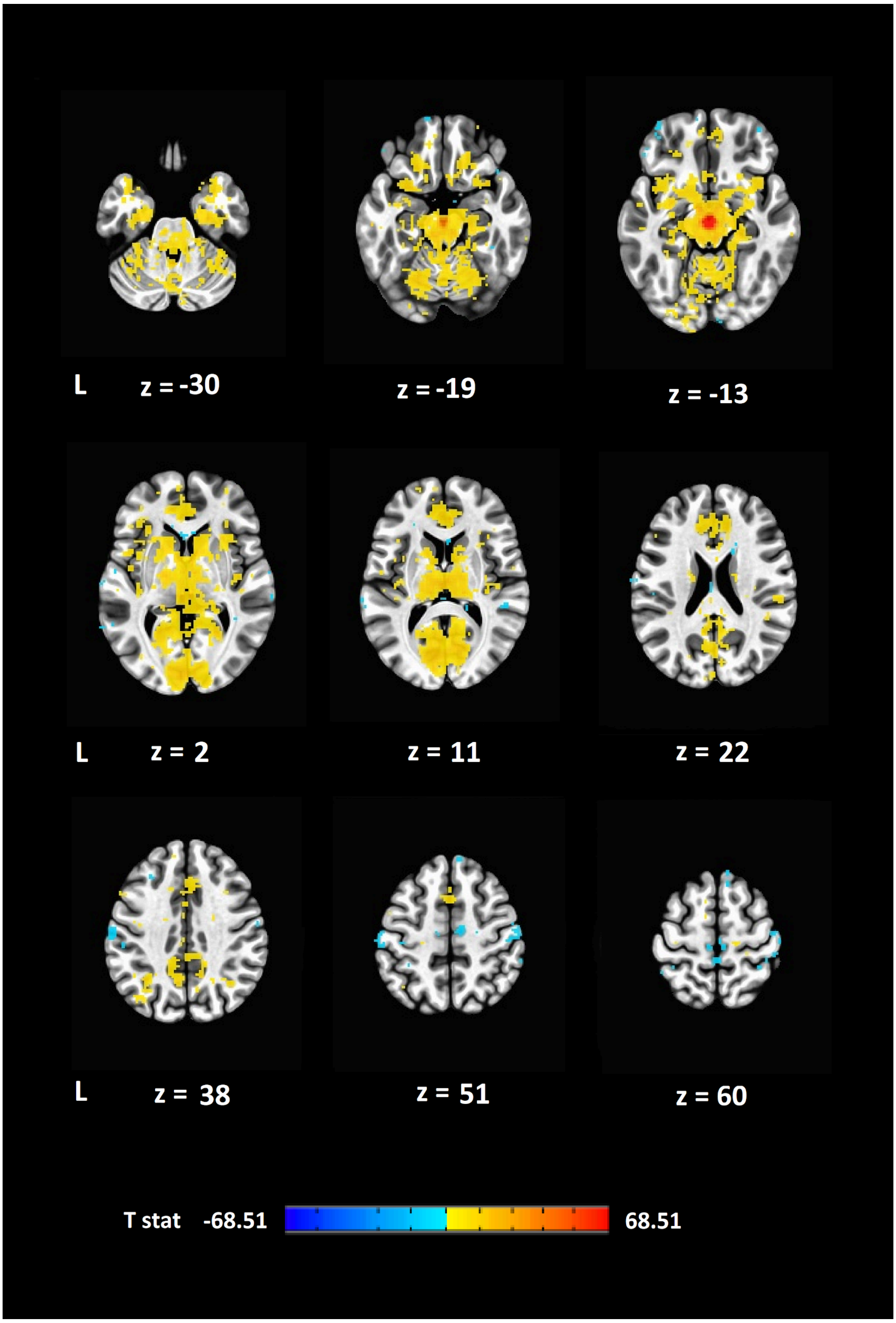


**Supplementary Figure 2.** Resting-state functional connectivity of ventral tegmental area (VTA). Warm and cool colors represent, respectively, positive and negative correlations.

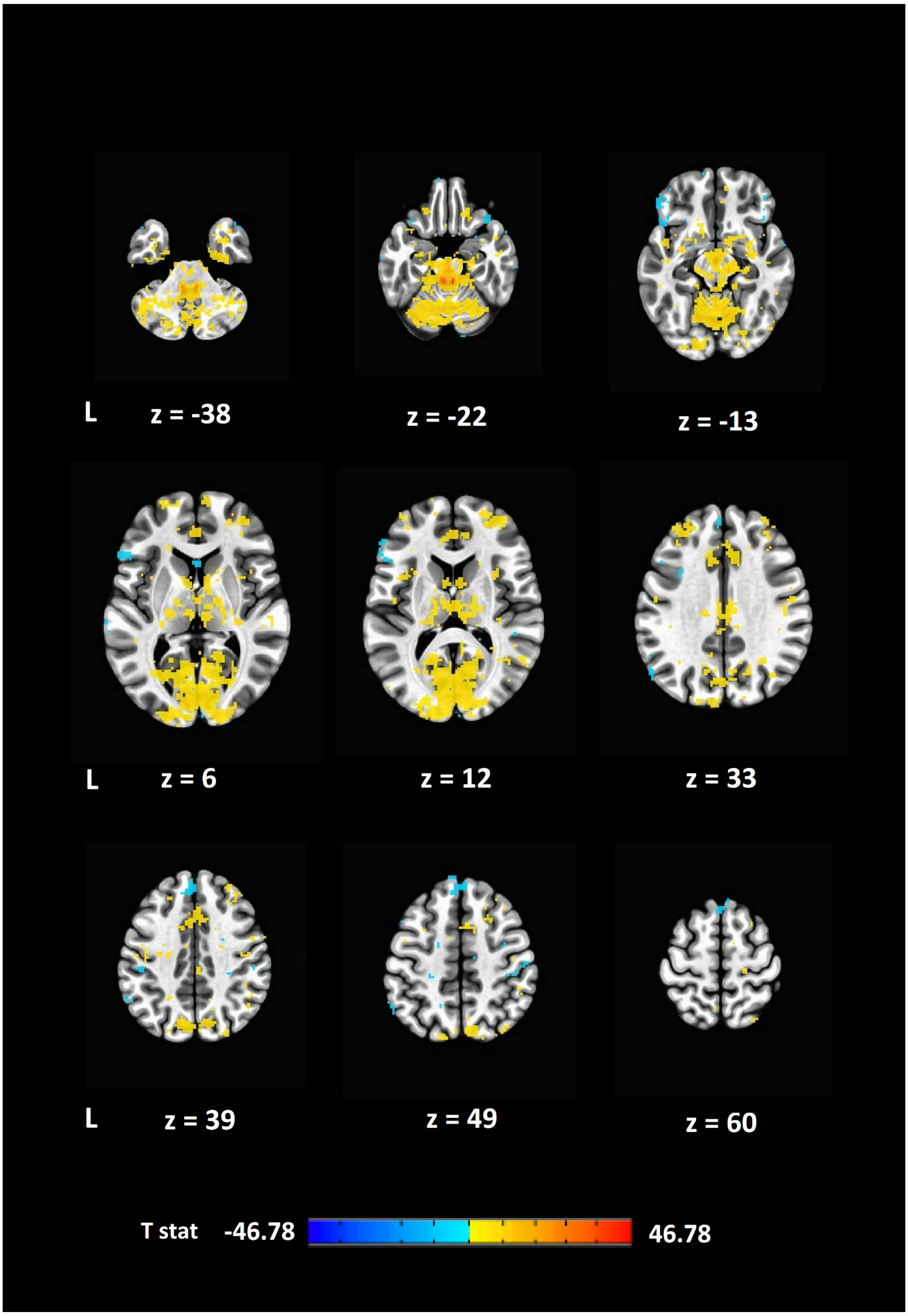


**Supplementary Figure 3.** Resting-state functional connectivity of locus coeruleus (LC). Warm and cool colors represent, respectively, positive and negative correlations.

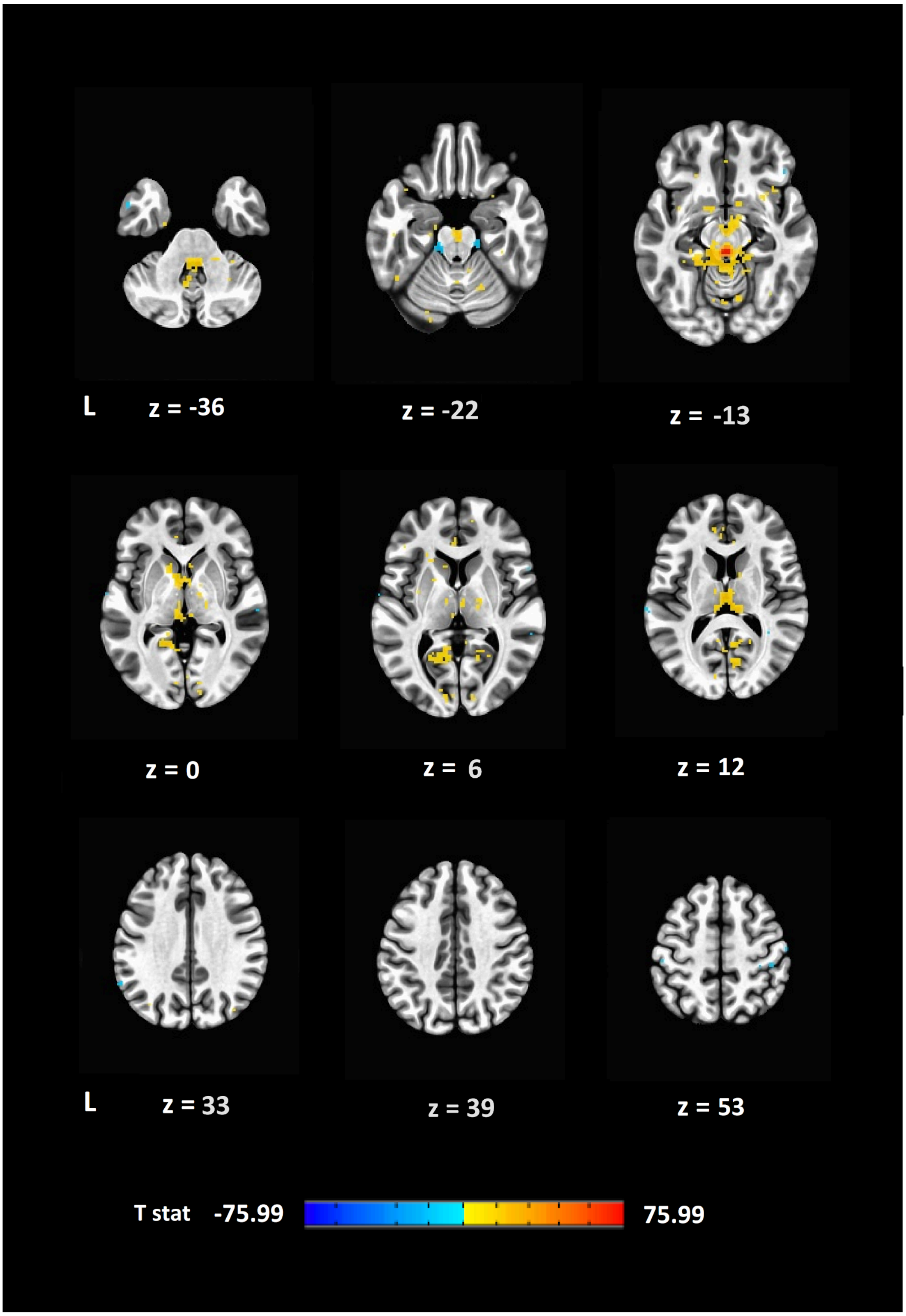


**Supplementary Figure 4.** Resting-state functional connectivity of dorsal raphe (DR). Warm and cool colors represent, respectively, positive and negative correlations.

### References


Abraham, Alexandre, Fabian Pedregosa, Michael Eickenberg, Philippe Gervais, Andreas Mueller, Jean Kossaifi, Alexandre Gramfort, Bertrand Thirion, and Gael Varoquaux. 2014. “Machine Learning for Neuroimaging with Scikit-Learn.” Frontiers in Neuroinformatics 8. https://doi.org/10.3389/fninf.2014.00014.

Andersson, Jesper L. R., Stefan Skare, and John Ashburner. 2003. “How to Correct Susceptibility Distortions in Spin-Echo Echo-Planar Images: Application to Diffusion Tensor Imaging.” NeuroImage 20 (2): 870–88. https://doi.org/10.1016/S1053-8119(03)00336-7.

Avants, B. B., C. L. Epstein, M. Grossman, and J. C. Gee. 2008. “Symmetric Diffeomorphic Image Registration with Cross-Correlation: Evaluating Automated Labeling of Elderly and Neurodegenerative Brain.” Medical Image Analysis 12 (1): 26–41. https://doi.org/10.1016/j.media.2007.06.004.

Behzadi, Yashar, Khaled Restom, Joy Liau, and Thomas T. Liu. 2007. “A Component Based Noise Correction Method (CompCor) for BOLD and Perfusion Based fMRI.” NeuroImage 37 (1): 90–101. https://doi.org/10.1016/j.neuroimage.2007.04.042.

Cox, Robert W., and James S. Hyde. 1997. “Software Tools for Analysis and Visualization of fMRI Data.” NMR in Biomedicine 10 (4-5): 171–78. https://doi.org/10.1002/(SICI)1099-1492(199706/08)10:4/5<171::AID-NBM453>3.0.CO;2-L.

Esteban, Oscar, Ross Blair, Christopher J. Markiewicz, Shoshana L. Berleant, Craig Moodie, Feilong Ma, Ayse Ilkay Isik, et al. 2018. “fMRIPrep 23.0.0.” Software. https://doi.org/10.5281/zenodo.852659.

Esteban, Oscar, Christopher Markiewicz, Ross W Blair, Craig Moodie, Ayse Ilkay Isik, Asier Erramuzpe Aliaga, James Kent, et al. 2018. “fMRIPrep: A Robust Preprocessing Pipeline for Functional MRI.” Nature Methods. https://doi.org/10.1038/s41592-018-0235-4.

Fonov, VS, AC Evans, RC McKinstry, CR Almli, and DL Collins. 2009. “Unbiased Nonlinear Average Age-Appropriate Brain Templates from Birth to Adulthood.” NeuroImage 47, Supplement 1: S102. https://doi.org/10.1016/S1053-8119(09)70884-5.

Gorgolewski, K., C. D. Burns, C. Madison, D. Clark, Y. O. Halchenko, M. L. Waskom, and S. Ghosh. 2011. “Nipype: A Flexible, Lightweight and Extensible Neuroimaging Data Processing Framework in Python.” Frontiers in Neuroinformatics 5: 13. https://doi.org/10.3389/fninf.2011.00013.

Gorgolewski, Krzysztof J., Oscar Esteban, Christopher J. Markiewicz, Erik Ziegler, David Gage Ellis, Michael Philipp Notter, Dorota Jarecka, et al. 2018. “Nipype.” Software. https://doi.org/10.5281/zenodo.596855.

Greve, Douglas N, and Bruce Fischl. 2009. “Accurate and Robust Brain Image Alignment Using Boundary-Based Registration.” NeuroImage 48 (1): 63–72. https://doi.org/10.1016/j.neuroimage.2009.06.060.

Jenkinson, Mark, Peter Bannister, Michael Brady, and Stephen Smith. 2002. “Improved Optimization for the Robust and Accurate Linear Registration and Motion Correction of Brain Images.” NeuroImage 17 (2): 825–41. https://doi.org/10.1006/nimg.2002.1132.

Jenkinson, Mark, and Stephen Smith. 2001. “A Global Optimisation Method for Robust Affine Registration of Brain Images.” Medical Image Analysis 5 (2): 143–56. https://doi.org/10.1016/S1361-8415(01)00036-6.

Lanczos, C. 1964. “Evaluation of Noisy Data.” Journal of the Society for Industrial and Applied Mathematics Series B Numerical Analysis 1 (1): 76–85. https://doi.org/10.1137/0701007.

Patriat, Rémi, Richard C. Reynolds, and Rasmus M. Birn. 2017. “An Improved Model of Motion-Related Signal Changes in fMRI.” NeuroImage 144, Part A (January): 74–82. https://doi.org/10.1016/j.neuroimage.2016.08.051.

Power, Jonathan D., Anish Mitra, Timothy O. Laumann, Abraham Z. Snyder, Bradley L. Schlaggar, and Steven E. Petersen. 2014. “Methods to Detect, Characterize, and Remove Motion Artifact in Resting State fMRI.” NeuroImage 84 (Supplement C): 320–41. https://doi.org/10.1016/j.neuroimage.2013.08.048.

Satterthwaite, Theodore D., Mark A. Elliott, Raphael T. Gerraty, Kosha Ruparel, James Loughead, Monica E. Calkins, Simon B. Eickhoff, et al. 2013. “An improved framework for confound regression and filtering for control of motion artifact in the preprocessing of resting-state functional connectivity data.” NeuroImage 64 (1): 240–56. https://doi.org/10.1016/j.neuroimage.2012.08.052.

Tustison, N. J., B. B. Avants, P. A. Cook, Y. Zheng, A. Egan, P. A. Yushkevich, and J. C. Gee. 2010. “N4itk: Improved N3 Bias Correction.” IEEE Transactions on Medical Imaging 29 (6): 1310–20. https://doi.org/10.1109/TMI.2010.2046908.
Zhang, Y., M. Brady, and S. Smith. 2001. “Segmentation of Brain MR Images Through a Hidden Markov Random Field Model and the Expectation-Maximization Algorithm.” IEEE Transactions on Medical Imaging 20 (1): 45–57. https://doi.org/10.1109/42.906424.
